# Revealing the antibiotic resistome in the global hadal trenches by large-scale cultivation and metagenomics

**DOI:** 10.64898/2025.12.11.692252

**Authors:** Xiang Cheng, Xiaojie Qiu, Naijia Shen, Hui Zhou, Mengjie Zhang, Zhe-Xue Quan, Guoqing Zhang, Jiasong Fang, Xuan Li, Guo-Ping Zhao, Pei Hao, Ping Chen

## Abstract

The global hadal trenches that reside more than 6000 meters below the sea level, represent one of Earth’s most remote ecosystems, potentially harboring a microbial reservoir for antibiotic resistant bacteria (ARB) and antibiotic resistance genes (ARGs). However, the global hadal environments remain the least investigated ecosystem for ARB/ARG compositions and distribution. This study took three-fold approaches to investigate the ARB/ARGs in the global hadal trench sediments, i.e., large-scale cultivation to isolate and verify ARB/ARGs from Challenger Deep sediments, metagenomic analysis of sediment samples from Challenger Deep over a six-year period, and comparative study of resistomes across six hadal trenches through metagenomic analysis. The Challenger Deep sediments had a consistent average ARG abundance of 0.02 copy/cell over a six-year period, whereas some ARG subtypes fluctuated and showed distinct temporal dynamics. Large-scale screening of ARB via microbial cultivation using 13 classes of antibiotics led to enriching distinct ARB, for which eleven genera were identified by metagenomic sequencing as major host for ARGs. The global hadal trench sediments showed a variable ARG abundance, which was significantly lower than those from environments in proximity to human activities, representing a relatively pristine reservoir. Phylogenetic analysis suggested the hadal ARGs underwent niche-specific selections and formed hadal trench-specific clusters that diverged from human-associated lineages, which can be outward transported when disturbed. The study established the hadal trench sediments, as a relatively pristine reservoir of natural ARGs, providing a crucial environmental baseline for assessing their impacts on the ecosystems and public health risks.

## Introduction

Antibiotic resistance genes (ARGs) are regarded as emerging environmental pollutants and a critical challenge to global public health [1]. Although ARGs have been documented in a wide range of ecosystems, the marine environment is increasingly recognized as an important reservoir of ARGs [2]. Among marine habitats, hadal trenches - the deepest and most remote zones of the ocean - remain largely unexplored in terms of their ARG diversity, distribution, and ecological significance. As components of a globally connected ecosystem, these extreme environments may play a crucial role in the biogeography and dissemination of resistance traits. The hadal zone harbors a rich diversity of uncharacterized microbial lineages [3, 4], suggesting the potential existence of novel resistance mechanisms. Moreover, profiling ARGs in these pristine ecosystems provides a baseline for assessing anthropogenic impacts on environmental resistome and may provide early warnings of ecological and public health risks. Current knowledge of hadal ARGs is limited to a few metagenomic studies, with examples of single-trench ARG analysis on the metagenomics data from the Yap trench and Mariana trench [5–7]. Although such culture-independent methods offer valuable insights, they cannot confirm phenotypic resistance or host association. The origins of ARGs in the extreme environments are drawing increasing attention in the research community. However, studies on ARGs and their origins in the global hadal zones is scarce, representing a significant gap in our understanding of the hadal zone resistome. To address these questions, the present study adopted a three-fold strategy to systematically investigate the compositions and distribution of antibiotic-resistant bacteria (ARB) and ARGs in the global hadal sediment environments. First, we conducted a large-scale cultivation to isolate and verify ARB/ARGs from the Challenger Deep sediment samples using 13 classes of antibiotics. Second, to track temporal changes in the hadal resistome, we analyzed metagenomic data from Challenger Deep sediments collected over a six-year period (2016-2021). Third, to systematically investigate the global distribution, composition and origins of ARGs in these extreme and remote ecosystems, we conducted a comparative metagenomic analysis of the resistome across six hadal trenches.

## Results and discussion

We first investigated the antibiotic resistome in the Challenger Deep of the Mariana Trench, the deepest hadal environment on earth. Metagenomic data for six continuous years, from 2016 to 2021, were collected (Table S1) for the sediment samples from the Challenger Deep (water depth of greater than 10000 m). The results indicated that the normalized abundance of total ARGs in the Challenger Deep sediments remained at consistent average levels (~0.02 copy/cell) over the six-year study period (ANOVA, *p* > 0.05) (Fig. 1a and Table S2). Although slightly elevated in 2020 and 2021 compared to the preceding four years, this increase was not statistically significant. A total of 24 ARG types were identified for them (Fig. 1b and Table S3), while 14 of them were commonly detected throughout the six-year period.

**Fig. 1.**
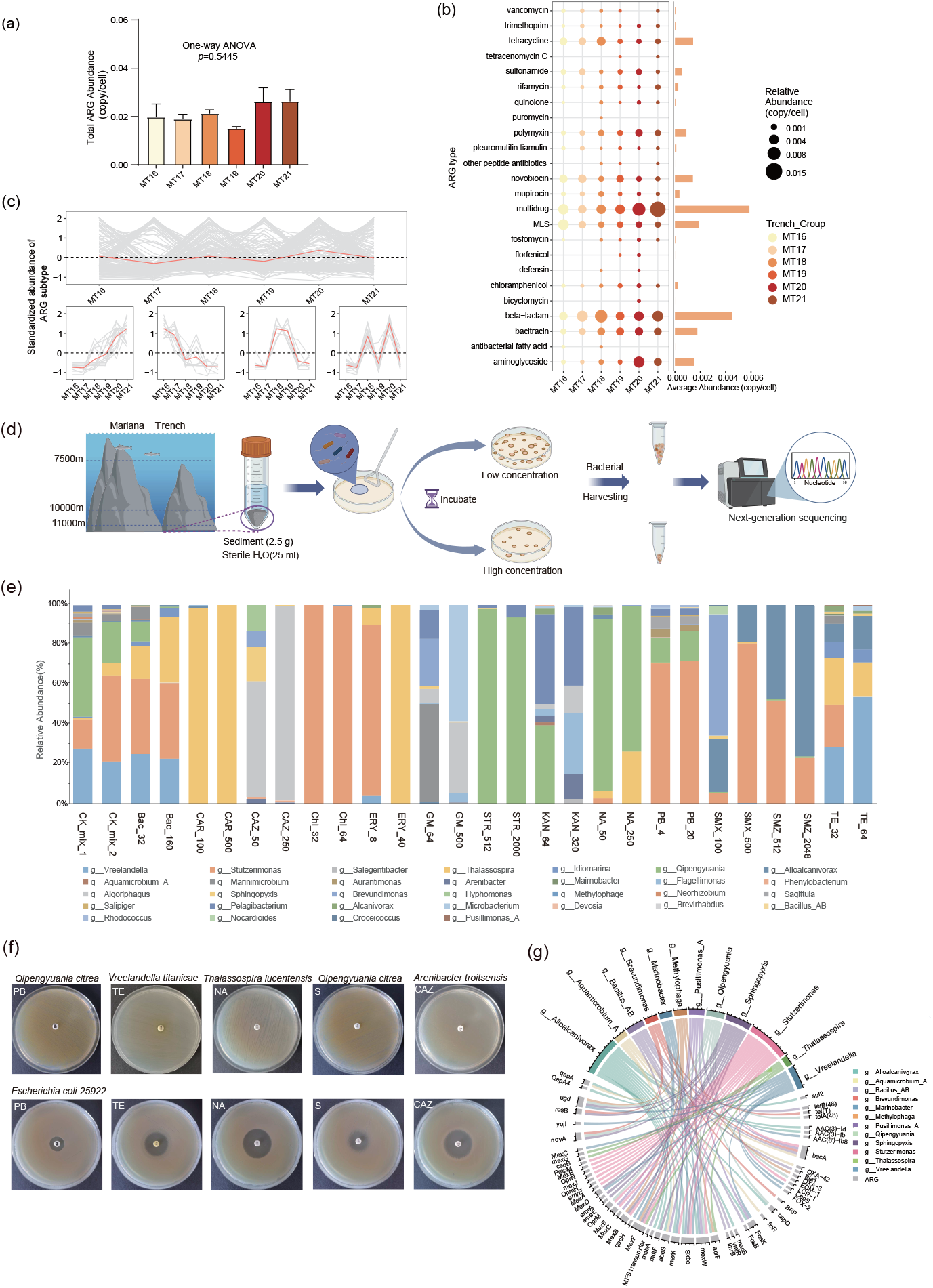
Comprehensive analysis of the antibiotic resistome and antibiotic-resistant bacteria in Mariana Trench Challenger Deep sediments. **(a)** The normalized abundance of the total ARGs (copy/cell) in Mariana Trench Challenger Deep sediments over the six-year (2016-2021) study period. Each error bar corresponds to the SEM and data were showed as mean + SEM. One-way ANOVA test was performed to obtain the *p* value. **(b)** Abundances and types of the ARGs detected in Mariana Trench Challenger Deep sediments over the six-year period. MLS: macrolide, lincosamide, and streptogramin. **(c)** Spatio-temporal variation of ARG subtype abundance in Mariana Trench Challenger Deep sediments from 2016 to 2021. The abundance was standardized with the “standardise” function in the R package Mfuzz (version 2.62.0). The top panel showed the fluctuation of all ARG subtypes over the six-year study period, and the bottom panels showed four distinct temporal dynamics of some small subsets. **(d)** Flowchart of antibiotic-resistant bacteria screening from hadal sediments. Antibiotic-resistant bacteria were isolated from Mariana Trench sediments using selective agar plates supplemented with a gradient of antibiotics. Detailed information on the antibiotics and the concentrations employed in this study is described in Table S8. The chart was created with the help of Biorender (https://Biorender.com). **(e)** Composition and relative abundance of antibiotic-resistant bacteria at genus level recovered from treatment with 13 distinct antibiotics. CAR_100, CAR is the abbreviation for the antibiotic and 100 indicates its concentration; CK_mix_1 and CK_mix_2 are two replicate experiments without any antibiotic. **(f)** Kirby–Bauer disk diffusion assay for drug resistance verification. Escherichia coli ATCC 25922 was included as the quality-control (QC) strain. PB: polymyxin B; TE: tetracycline; NA: nalidixic acid; S: streptomycin; CAZ: ceftazidime. **(g)** Chord diagram analysis revealing the mapping relationship between the ARG-carrying MAGs (at the genus level) and the carried ARGs.

Multidrug, beta-lactam, bacitracin, and macrolide-lincosamide-streptogramin (MLS) were the most frequently detected ARG types. Looking more closely, on average 82 ARG subtypes were found in the sediments, among which *PNGM-1, bacA, novA, macB* and *msbA* were consistently abundant over the six-year period (Table S4). While the overall ARG reservoir remains relatively stable at the community level (Fig. 1a), individual ARG subtypes undergo fluctuation in their abundance (Fig. 1c top panel), among which some small subsets showed distinct temporal dynamics (Fig. 1c bottom panel). For instance, *bacA* and *dfrA*16, displayed a progressive increase in relative abundance from 2016 to 2021. Conversely, genes such as *novA, rphB*, and *tetB(P)*, showed a significant decline over the same period. Other ARGs, such as *mdtF, mdtN* and *mdtP*, exhibited periodic oscillations between 2016 and 2021. Analysis of ARGs in Challenger Deep sediments identified efflux pumps as the primary resistance mechanism (Table S5 and Fig. S1a online), predominantly multidrug resistance (MDR) types (Table S6). This finding agrees with previous studies suggesting MDR efflux pumps are evolutionarily ancient and highly conserved [8]. Although ARGs generally pose a threat to public health, application of an omics-based risk framework (Ranks I-IV) [9] showed that high-risk ARGs (Ranks I-II) accounted for only ~2.7% of the local resistome over a six-year period (Table S7 and Fig. S1b online). This proportion is markedly lower than those reported for wastewater (67.0%) and deep-sea cold seep microbiomes (11.26%) [10], indicating that the Challenger Deep resistome presents a very low potential risk to public health.

Current approaches for ARG study primarily rely on sequencing-based techniques, which did not verify whether the potential ARGs are functionally active within living hosts. To bridge this gap and functionally characterize the ARGs/ARB in the Challenger Deep sediments, we conducted large-scale screening of ARB via microbial culture experiments using 13 classes of antibiotics, with two concentrations for each screening antibiotic (Table S8). Growing bacterial colonies were collected from the plates and subjected to metagenomic sequencing for each plate separately (Fig. 1d). Metagenome assembly and binning yielded 189 high-quality MAGs (Table S9). Taxonomic classification assigned them to 4 bacterial phyla (Pseudomonadota, Bacteroidota, Actinomycetota, and Bacillota) and 5 classes (Alphaproteobacteria, Gammaproteobacteria, Bacteroidia, Actinobacteria, and Bacilli), encompassing 15 orders, 21 families, and 32 genera. Metagenomic analysis revealed distinct taxonomic compositions and abundance patterns among the colonies from different antibiotic treatments (Fig. 1e and Table S10). For instance, chloramphenicol plates were dominated by genera such as *Stutzerimonas*, while streptomycin and carbenicillin plates were dominated by *Qipengyuania* and *Thalassospira*, respectively. Furthermore, similar ARBs were consistently recovered across different concentrations of the same antibiotic (Fig. 1e). These results indicated that different bacterial genera possessed distinct antibiotic resistance profiles. Moreover, NMDS analysis based on Bray-Curtis dissimilarity demonstrated significant, antibiotic-driven clustering of microbial communities (PERMANOVA, *p* = 0.001, *R*^2^ = 0.846) (Fig. S2 online), affirming deterministic selection rather than stochastic drift. To validate the antibiotic resistance of isolates from screening plates, we performed disk diffusion assays on a subset of colonies. Compared to the reference strain *Escherichia coli* ATCC 25922 that exhibited clear antibiotic inhibition zones (Fig. 1f bottom), isolates from PB-, TE-, CAZ-, STR-, and NA-plates showed no inhibition and grew on the corresponding antibiotic-containing plates (Fig. 1f top). Notably, *Qipengyuania citrea* - isolated from both PB- and STR-plates - exhibited a multidrug-resistant (MDR) phenotype, demonstrating concurrent resistance to polymyxin and streptomycin. These results confirm the true resistance of the isolates, delineate species-specific resistance profiles. We further characterized the ARB selected on the antibiotic plates and their associated ARGs. Specifically, we identified 1504 ARG-containing ORFs, classified into 18 types and 110 subtypes (Table S11). For instance, in carbenicillin (CAR)-containing plate, we identified CAR-specific resistance genes, including the the β-lactamase gene *LCR-1* and the efflux pump system *MexAB-OprM*. In polymyxin B (PB)-containing plate, in addition to the broad-spectrum efflux system *MexAB-OprM*, we detected PB-specific efflux genes such as *RosAB*, as well as the *ugd* gene, which modifies lipid A to reduce PB binding affinity. In chloramphenicol (Chl)-containing plate, broad-spectrum efflux pump genes including *MexAB-OprM* and *MexEF-OprN* were identified (Table S11). To identify the bacterial hosts carrying these resistance genes, we analyzed the 189 high-quality MAGs, which represented 32 distinct genera. Our analysis showed that all 32 genera carried ARGs. Eleven of these genera were identified as major carriers, such as *Marinobacter, Qipengyuania*, and *Stutzerimonas* (Fig. 1g), with each harboring at least five different ARGs, suggesting that they constitute key resistance reservoirs in the hadal zone. Notably, we identified a strain of *Stutzerimonas chloritidismutans* that grew on multiple antibiotics.

Genomic analysis revealed the molecular basis for its multidrug resistance: the efflux pump genes *mexAB-oprM* (conferring broad-spectrum resistance) and *mexEF-oprN* (linked to chloramphenicol resistance), along with the bacitracin resistance gene *bacA*. This genotype-phenotype correlation was consistently observed. The presence of *sul2, acrF*, and *oqxB* in *Alloalcanivorax venustensis* correlated with its growth on carbenicillin (CAR), sulfamethoxazole (SMX), sulfamethazine (SMZ), and tetracycline (TE). Similarly, *rosA* and *AAC(3)-IIIb* genes in *Pelagibacterium* sp. corresponded to resistance to polymyxin B (PB), streptomycin (STR), and kanamycin (KAN) (Table S12 and Fig. S3 online). In summary, the analysis of MAGs from cultured isolates revealed a clear genotype-phenotype link, confirming that hadal bacteria harbor diverse ARGs and providing key insights into the dissemination and evolution of antibiotic resistance in this extreme environment.

Studies of ARB/ARGs in the global hadal trench environments are scarce. We performed the first investigation of ARGs in hadal trench sediments globally. Metagenomic data from six hadal trenches, Mariana Trench (MT), Diamantina Trench (DT), Massau Trench (MST), New Britain Trench (NBT), Yap (YT), and Kermadec Trench (KT), were analyzed (Table S1). The average ARG abundance across trenches was approximately 0.02 copy/cell (Table S13), with DT and MST showing lower, albeit statistically insignificant, abundances (Fig. 2a). In contrast, human-influenced environments such as rivers, lakes, and estuarine/coastal areas exhibited ARG abundances of 0.3-1.5 copy/cell [11], at least an order of magnitude higher. These results underscore the relatively pristine state of hadal sediments and reinforce the role of human activity in driving ARG spread in nature. A total of 25 ARG types were identified, 15 of which were common to all trenches (Fig. 2b and Table S14); among these, beta-lactam, multidrug, bacitracin, and MLS were the dominant types. Subtypes such as *bacA, PNGM-1, macB*, and *msbA*, were with high abundance and widespread across the hadal trench, whereas human-associated subtypes, such as *TEM-1, sul1*, and *sul2*, were detected only in a portion of global hadal trenches and at relatively low abundance (Table S15 and Fig. S4 online). Unlike human-impacted coastal/estuarine regions where sulfonamide (e.g., *sul1, sul2*) and tetracycline (e.g., *tetA, tetW*) resistance genes prevail [12, 13], hadal environments are populated by functional resistance genes related to extreme-environment adaptation, such as multidrug efflux pumps (e.g., *msbA* and *RanA*), and intrinsic resistance genes (e.g. *bacA)*. To assess ARG diversity of global hadal sediments, our analysis of six trenches revealed an average richness of 70 ARG subtypes per trench (Fig. S5a online), which is lower than that reported for estuarine/coastal sediments (~100 subtypes) [14], freshwater sediments (~200 subtypes) [15], and untreated sewage (~400 subtypes) [15]. While richness did not differ significantly among trenches, diversity indices indicated uneven ARG distribution in KT and NBT (Fig. S5b,c online). Efflux pumps and enzymatic inactivation were found to be the primary resistance mechanisms in all trenches (Fig. 2c and Table S16). In agreement with what we found in the MT sediments earlier, the vast majority (>90%) of ARGs in all trenches were classified as low-risk (“Rank IV”; Fig. 2d and Table S17). The high-risk ARGs (Ranks I-II) were rarely detected across the trenches, suggesting a minimal public health risk.

**Fig. 2.**
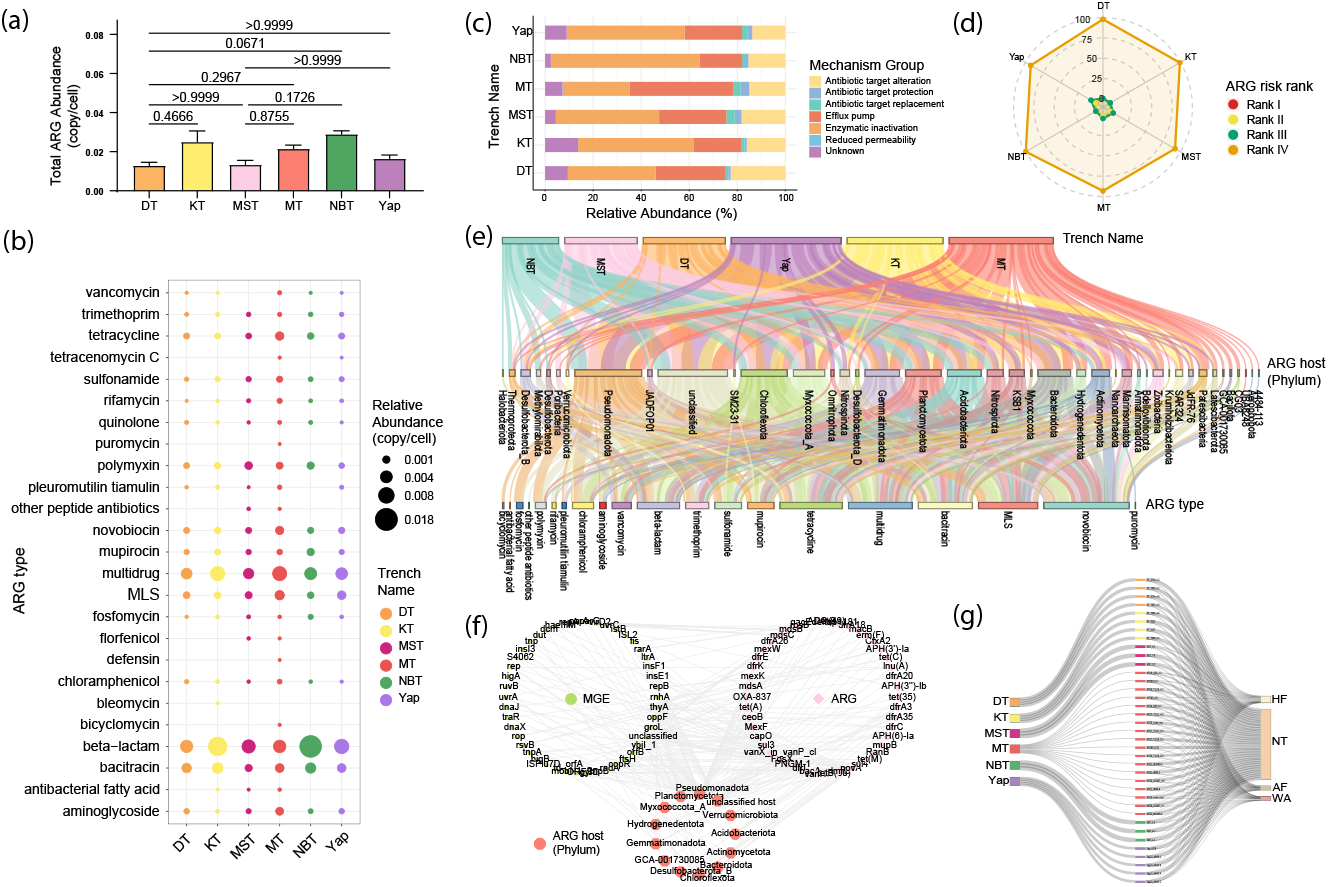
The antibiotic resistome in the sediment environments of the global hadal trenches. **(a)** The normalized abundance of the total ARGs (copy/cell) in the six hadal trench sediments. Each error bar corresponds to the SEM and data were showed as mean + SEM. Dunn’s multiple comparison test with Bonferroni correction was performed to obtain the *p* value. **(b)** Abundances and types of the ARGs detected in the six hadal trench sediments. **(c)** Composition of ARG resistance mechanisms in different hadal trenches. **(d)** Radar plot showing the relative proportion of different risk ranks of ARGs distributed across the six hadal trenches. **(e)** The mapping relationship between the ARGs (at the type level, 3rd row) and their potential hosts (at the phylum level, 2nd row) in the six hadal trench sediments. **(f)** Visualization of the ARG-MGE-host association network. Ellipse, Diamond and Octagon denote MGE, ARG and microbial hosts, respectively. Association is summarized among MGE element, ARG subtype and a microbe phylum, represented by a linked grey line. **(g)** Sankey diagram illustrating the relative contribution of various sources (i.e. AF, HF, NT and WA) to the ARGs in the six hadal trench predected by SourceTracker. NT: natural environment; AF: animal feces; HF: human feces; WA: wastewater treatment plant; MT: Mariana Trench; NBT: New Britain Trench; Yap: Yap Trench; MST: Massau Trench; KT: Kermadec Trench; MLS: macrolide, lincosamide, and streptogramin.

Given the profile of ARG distribution in global hadal trench sediments, we next investigated what their hosts are and how ARGs disseminate in the harsh environments. Metagenomic assembly of the global hadal trench sediments yielded 9,852,945 contigs (>1 kb) (Table S18), of which 0.04% carried ARGs. These contigs were taxonomically assigned to 40 phyla and 270 genera (Table S19), with dominant ARG hosts including Pseudomonadota, Chloroflexota, and Gemmatimonadota (Fig. 2e,). Archaeal carriers (e.g., Thermoplasmatota, Nanoarchaeota) were also detected. A notable fraction of ARG-carrying contigs remained unclassified (Fig. 2e), suggesting novel microbial lineages in hadal settings. A total of 177 ARG subtypes were identified from the carrying contigs, which could enable resistance to 20 types of antibiotics (Fig. 2e and Table S19). To explore how ARGs disseminate in hadal trenches, we analyzed the mobile genetic elements (MGEs) co-occurring with ARGs on the same contigs. Our analysis identified 200 ARG-MGE pairs across 13 phyla, primarily in Pseudomonadota and Chloroflexota (Fig. 2f and Table S20). MGE composition was similar across trenches (Fig. S6 online), with integration/excision and phage-related elements being most common. Only 4.28% of ARG-carrying contigs were linked to MGEs, indicating limited horizontal transfer in hadal sediments. To look into ARGs presence at a whole genome level, we constructed MAGs. Among the recovered 1,344 MAGs (Table S21), 18.7% contained at least one ARG (Table S22). This prevalence is substantially lower than that in environments impacted by human activity, such as mining areas (85%) [15], wastewater treatment systems (57%) [16], and drinking water reservoirs (40%) [11], as well as in natural/engineered subsurface systems, including Tibetan Plateau groundwater (84.6%) [17] and anchialine caves (55%) [18]. Most ARG-carrying MAGs belonged to Pseudomonadota, Chloroflexota, and Actinomycetota (Fig. S7). Only 5.58% of these MAGs co-harbored MGEs (Fig. S7), a proportion markedly lower than in other environments (e.g., 37% in anchialine caves [18]). Consistent with the aforementioned results at the ARG-carrying contigs level, the low prevalence of MGEs supports the notion that vertical inheritance, rather than horizontal transfer, constitutes the primary mode of ARG dissemination in pristine environments such as the hadal sediments.

To understand the origins of ARGs in global hadal trench sediments, we applied Bayesian source-tracking models trained on 594 metagenomic datasets from four ecotypes - human feces (HF), animal feces (AF), wastewater treatment plants (WA), and natural environments (NT) - spanning 12 niches and 6 continents (Table S23) to predict their sources. Results identified NT as the dominant source of hadal ARGs, contributing approximately 70% overall (Fig. 2g and Table S24). NT was also the primary source for the most abundant ARG types such as multidrug, MLS, and beta-lactam (Table S25 and Fig. S8 online). Phylogenetic analyses further revealed the natural and niche-specific evolution of hadal ARGs. For instance, hadal *fosX* sequences clustered within Clade I, closely related to sequences derived from natural environments like the North Pacific and hydrothermal vents, while Clade II - containing pathogens such as *Listeria monocytogenes* - was distantly related (Fig. S9a online). Similarly, *sul2* and *arnA* genes from hadal sediments grouped with homologs from natural habitats including hot springs, marine systems, and deep-sea sediments, showing clear divergence from human-associated lineages (Fig. S9b,c online). In the case of the efflux pump gene *abeS* (Fig. S9d online), hadal sequences formed a unique clade (Clade I), separate from all other sources, suggesting these genes represent ancient, niche-adapted lineages that evolved independently in hadal settings. Together, our source-tracking and phylogenetic results indicated the pristine nature of hadal sediments, where hadal ARGs are primarily natural in origin and have undergone habitat-specific selection, forming distinct genetic reservoirs that may be outward transported if disturbed.

## Conclusions

In conclusion, our study presents the first comprehensive analysis of ARB/ARGs in the sediments of global hadal trenches, the deepest and most remote marine ecosystems on Earth. Through a three-fold approach, i.e. large-scale cultivation (using 13 classes of antibiotics) to isolate and verify ARB/ARGs from Challenger Deep sediments, metagenomic analysis of continuously sampled Challenger Deep sediments over a six-year period, and comparative study of resistome across six hadal trenches via metagenomics, we revealed for the first time, the distribution and compositions of ARB/ARGs in the global trench environments, and also the possible origins of these ARB/ARGs. We found that the Challenger Deep sediments had a consistent average abundance of 0.02 ARG copy/cell over a six-year period, whereas many ARG subtypes fluctuated, subsets of which showed distinct temporal dynamics.

By large-scale screening of ARB via microbial cultivation using 13 classes of antibiotics, we found that bacterial communities were structured differently in response to various antibiotic classes, with each antibiotic enriching distinct genera of ARB. Among the 32 genera of ARB enriched on the antibiotic plates, eleven were identified as major hosts of ARGs. The comparative analysis of global hadal sediments showed a variable ARG abundance among them, which were significantly lower than those from environments heavily influenced by human activities. The results indicated that hadal environments represent a relatively pristine reservoir. Interestingly, the hadal trench sediments were different in ARG types between them, whereas subtypes that were often found in vicinity close to human activities, were detected only in a portion of global hadal trenches and at relatively low abundance. Across global hadal trenches, a total of 177 ARG subtypes were identified via metagenomic assembly, conferring potential resistance to 20 antibiotic classes. Their harboring host bacteria belong to 40 phyla and 270 genera. We also found that MGEs had a low prevalence, which supports the notion that vertical inheritance, rather than horizontal gene transfer, constitutes the primary mode of ARG dissemination in the pristine environment. Source tracking analysis indicated that NT may be the major source of hadal ARGs, whose relative contribution account for around 70%. Further phylogenetic evidence suggested that ARGs underwent niche-specific selections in the hadal trench sediment environments, and formed hadal trench-specific clusters of ARGs which can become sources of outward transport when disturbed. The study established the hadal trench sediments as a pristine reservoir of natural ARGs, providing a crucial environmental baseline for assessing the impact of anthropogenic activity on the global spread of antibiotic resistance. Despite these limitations, this study underscores the value of conserving these remote ecosystems as natural laboratories for understanding the pre-anthropogenic evolution of antibiotic resistance.

## Data availability

The metagenomic sequencing data for hadal trench sediment samples and the genomic DNA data derived from culturomics-cultivated antibiotic-resistant bacterial colonies were deposited in the National Center for Biotechnology Information (NCBI) Sequence Read Archive (SRA) under BioProject accession numbers PRJNA1335898 (sediment samples) and PRJNA1335899 (culturomics-derived resistant bacteria), respectively.

## Acknowledgements

This work was supported in part by grants from the National Natural Science Foundation of China (92451303, 32200093), the China Postdoctoral Science Foundation (2022M713144), and the National Key R&D Program of China (2018YFC0310600, 2018YFA0900700). The authors thank the crews of the R/V “Shen Kuo”, R/V “Tan Suo Yi Hao”, and R/V “Zhang Jian” for cruise organization, technological supporting, and assistance on sampling.

## Author contributions

X.L., P.C., G.Z. and P.H. conceived the project. P.C., X.C. and X.L. wrote the manuscript. P.C., X.C., X.Q., N.S. and M.Z. conducted metagenomics experiments, analyzed data, and prepared the figures. X.C. performed antibiotic-resistant bacteria cultivation and isolation. J.F. and H.Z. collected hadal sediment samples. Z.Q. and G.Z. advised on the study.

## References

[1] Collaborators AR. Global burden of bacterial antimicrobial resistance in 2019: A systematic analysis. Lancet, 2022, 399: 629–655

[2] Xu N, Qiu D, Zhang Z, et al. A global atlas of marine antibiotic resistance genes and their expression. Water Res, 2023, 244: 120488

[3] Xiao X, Zhao W, Song Z, et al. Microbial ecosystems and ecological driving forces in the deepest ocean sediments. Cell, 2025, 188: 1363–1377

[4] Chen P, Zhou H, Huang Y, et al. Revealing the full biosphere structure and versatile metabolic functions in the deepest ocean sediment of the challenger deep. Genome Biol, 2021, 22: 207

[5] Su H, Wu C, Han P, et al. The microbiome and its association with antibiotic resistance genes in the hadal biosphere at the yap trench. J Hazard Mater, 2022, 439: 129543

[6] Sun J, Xie YG, Zhou H, et al. Distribution patterns and ecological risks of antibiotic resistance genes in the yap trench. Water Res, 2025, 281: 123589

[7] Yang H, Liu R, Liu H, et al. Evidence for long-term anthropogenic pollution: The hadal trench as a depository and indicator for dissemination of antibiotic resistance genes. Environ Sci Technol, 2021, 55: 15136–15148

[8] Martinez JL, Sánchez MB, Martínez-Solano L, et al. Functional role of bacterial multidrug efflux pumps in microbial natural ecosystems. FEMS Microbiol Rev, 2009, 33: 430–449

[9] Zhang AN, Gaston JM, Dai CL, et al. An omics-based framework for assessing the health risk of antimicrobial resistance genes. Nat Commun, 2021, 12: 4765

[10] Zhang T, Han Y, Peng Y, et al. The risk of pathogenicity and antibiotic resistance in deep-sea cold seep microorganisms. mSystems, 2025, 10: e0157124

[11] Wang S, Nie W, Gu Q, et al. Spread of antibiotic resistance genes in drinking water reservoirs: Insights from a deep metagenomic study using a curated database. Water Res, 2024, 256: 121572

[12] Leng Y, Xiao H, Li Z, et al. Tetracyclines, sulfonamides and quinolones and their corresponding resistance genes in coastal areas of beibu gulf, china. Sci Total Environ, 2020, 714: 136899

[13] Zhang Y, Lu J, Wu J, et al. Occurrence and distribution of antibiotic resistance genes in sediments in a semi-enclosed continental shelf sea. Sci Total Environ, 2020, 720: 137712

[14] Wu C, Zhang G, Zhang K, et al. Strong variation in sedimental antibiotic resistomes among urban rivers, estuaries and coastal oceans: Evidence from a river-connected coastal water ecosystem in northern china. J Environ Manage, 2023, 342: 118132

[15] Yi X, Liang JL, Su JQ, et al. Globally distributed mining-impacted environments are underexplored hotspots of multidrug resistance genes. Isme j, 2022, 16: 2099–2113

[16] Zhu C, Wu L, Ning D, et al. Global diversity and distribution of antibiotic resistance genes in human wastewater treatment systems. Nat Commun, 2025, 16: 4006

[17] Yang Y, Li H, Wang D, et al. Metagenomics of high-altitude groundwater reveal different health risks associated with antibiotic-resistant pathogens and bacterial resistome in the latitudinal gradient. Water Res, 2024, 262: 122032

[18] Vojvoda Zeljko T, Kajan K, Jalžić B, et al. Genome-centric metagenomes unveiling the hidden resistome in an anchialine cave. Environ Microbiome, 2024, 19: 67

