## Supplementary materials for "Revealing the antibiotic resistome in the global hadal trenches by large-scale cultivation and metagenomics"

­

Xiang Cheng^1, 10, 11^, Xiaojie Qiu^3^, Naijia Shen^2, 4, 10^, Hui Zhou^1^, Mengjie Zhang^1^, Zhe‑Xue Quan^5^, Guoqing Zhang^7^, Jiasong Fang^9^, Xuan Li^1, 10,*^, Guo‑Ping Zhao^5, 6, 7, 8,*^, Pei Hao^2, 4, 10,*^, and Ping Chen^2, 4, 11, *^

^1^Key Laboratory of Synthetic Biology, State Key Laboratory of Plant Trait Design, CAS Center for Excellence in Molecular Plant Sciences, Chinese Academy of Sciences, Shanghai, China

^2^Shanghai Institute of Materia Medica, Chinese Academy of Sciences, Shanghai, China

^3^School of Life Sciences, Henan University, Kaifeng, Henan, China

^4^Key Laboratory of Molecular Virology and Immunology, Shanghai Institute of Immunity and Infection, Chinese Academy of Sciences, Shanghai, China

^5^State Key Laboratory of Wetland Conservation and Restoration, School of Life Sciences, Fudan University, Shanghai, China

^6^State Key Laboratory of Genetics and Development of Complex Phenotypes, Fudan Microbiome Center, and School of Life Sciences, Fudan University, Shanghai, China.

^7^Bio-Med Big Data Center, Shanghai Institute of Nutrition and Health, Chinese Academy of Sciences, Shanghai, China

^8^School of Life Science, Hangzhou Institute for Advanced Study, University of Chinese Academy of Sciences, Hangzhou, China

^9^Shanghai Engineering Research Center of Hadal Science and Technology, College of Marine Sciences, Shanghai Ocean University, Shanghai, China

^10^University of Chinese Academy of Sciences, Beijing, China

^11^These authors contributed equally to this work

* Corresponding author:

Xuan Li, Pei Hao, Ping Chen,

Guo‑Ping Zhao

**Materials and Methods**

**Collection of metagenomic samples**

We collected metagenomic data from hadal sediment samples (depth > 6000 m) obtained from six hadal trenches: the Mariana Trench (MT), Diamantina Trench (DT), New Britain Trench (NBT), Yap Trench (Yap), Massau Trench (MST), and Kermadec Trench (KT). A total of 37 samples were analyzed, including eighteen from various stations and depth layers in the Mariana Trench collected over a six-year period (2016-2021), along with four from the Diamantina Trench, three from the New Britain Trench, five from the Yap Trench, three from the Massau Trench, and four from the Kermadec Trench. Sediment sampling was conducted at multiple sites across these trenches during different research cruises, with detailed sample information provided in Table S1. Metagenomic data for a subset of samples were made available from previous studies , and corresponding NCBI-SRA accession numbers are listed in Table S1. Metagenomic data newly generated in this study were obtained through library construction and sequencing, as detailed in the “DNA Extraction and Metagenomic Sequencing” section.

**DNA Extraction and Metagenomic Sequencing**

Genomic DNA was extracted from hadal trench sediment samples using the TIANamp Soil DNA Kit (centrifuge column format, catalog no. DP336; Tiangen Biotech Co., Ltd., Beijing, China) following the manufacturer’s protocols. For culturomics-derived resistant bacterial colonies (see “Isolation, identification, and antimicrobial susceptibility profiling of cultural antibiotic-resistant bacteria from Challenger Deep sediments”), genomic DNA was isolated via the CTAB method .

Illumina-compatible sequencing libraries were prepared using the Rapid Plus DNA Lib Prep Kit for Illumina (RK20208) (ABclonal, China). Qualified libraries were pooled and sequenced on the Illumina NovaSeq X Plus platform (Illumina, San Diego, CA, USA) with PE150 strategy in Novogene Bioinformatics Technology Co., Ltd (Beijing, China), according to effective library concentration and the data amount required.

**Quality Control, Assembly, and Binning of Metagenomic Sequencing Data**

Raw metagenomic reads were evaluated with FastQC (v0.12.1), and subsequently processed with Trimmomatic (v0.39) for adapter trimming and removal of low-quality sequences using the following parameters: ILLUMINACLIP:TruSeq3-PE-2.fa:2:30:10:8:true LEADING:3 TRAILING:3 SLIDINGWINDOW:4:15 HEADCROP:5 MINLEN:50 AVGQUAL:20. The resulting high-quality outputs were retained as clean reads for subsequent downstream analysis. Clean reads were then randomly subsampled using SeqKit to obtain comparable read counts across samples. The subsampled clean reads were subsequently merged using bbmap (v38.18) with parameters: rem k=60 extend2=50 ecct vstrict. Both the merged paired-end reads and the subsampled, quality-controlled reads were subsequently assembled into contigs using MEGAHIT (v1.2.9) with the assembly parameters reported for Challenger Deep sediments --k-list 21,29,39,59,79,99,119,127,139. The assembled contigs were selected for metagenomic binning using MetaWRAP (v1.3.2) . Briefly, initial binning was conducted with MetaBAT2, MaxBin2, and CONCOCT, leveraging their complementary strengths to generate a comprehensive set of draft genomes. The resulting bins from all three binning tools were subsequently integrated and refined through the MetaWRAP bin_refinement module with options: -c 50 -x 10. A cutoff of 50% completeness and 10% contamination was used to filter and obtain quality metagenome-assembled genomes (MAGs). The MAGs from the same hadal trench were further dereplicated using dRep v3.3.054 with the parameters ‘--P_ani 0.98 --S_ani 0.98 -nc 0.8’. The relative abundance of MAGs were quantified using the metaWRAP quant_bins module (v1.3.2).

**Reads-based detection and quantification of ARGs**

ARGs-OAP (v3.2.4) was used to characterize the antibiotic resistance gene (ARG) profiles based on the quality-controlled reads. The pipeline employs BLASTX to align putative ARG sequences against the Structured ARG Database (SARG v3). To account for differences in read length across hadal-trench datasets, we applied read-length–specific alignment thresholds. For the 150-bp datasets, we used an E-value of 1×10⁻⁷, identity ≥ 80% identity, and query coverage ≥ 75%. For the 100-bp dataset, we used an E-value of 1×10⁻⁷, identity≥85%, and query coverage ≥ 95%. For the 300-bp datasets, we used an E-value of 1×10⁻⁷, identity ≥ 75%, and query coverage ≥ 60%. ARG abundances were normalized to predicted cell counts and reported as copies per cell.

**Taxonomic profiling of microbial communities**

Taxonomic identification of the contigs was performed using CAT (v6.0) . The Genome Taxonomy Database (GTDB, release R220) was downloaded and formatted for CAT. The contigs were then aligned against this custom database, followed by taxonomic assignment using the “add_names” tool within CAT. GTDB-Tk (v2.4.0) based on the Genome Taxonomy Database (GTDB, R220) was used for taxonomic identification of MAGs.

**Metagenomic Assembly for Identification of ARG-like Open Reading Frames**

Open reading frames (ORFs) were predicted using Prodigal (v2.6.3) from assembled contigs. DIAMOND BLASTP (v2.16.0+) was then used to search these ORF sequences against the Structured ARG Database (SARG v3) with an E-value threshold of 1e^-7^. An ORF was identified as an ARG-like ORF if its best hit met both thresholds: similarity ≥ 60% and query coverage ≥ 60%.

**Isolation, identification, and antimicrobial susceptibility profiling of cultural antibiotic-resistant bacteria from Challenger Deep sediments**

The Challenger Deep sediments collected in 2018 (Table S1) were used to obtain cultural antibiotic-resistant bacteria. The screening process employed 13 distinct antibiotic agents, including: bacitracin, carbenicillin, gentamicin, chloramphenicol, kanamycin, polymyxin B, ceftazidime, streptomycin, sulfamethazine, sulfamethoxazole, erythromycin, nalidixic acid, and tetracycline. Each antibiotic was tested at two working concentrations, according to the 34th edition of the CLSI M100-Ed34 (2024), and two groups of bacteria cultured in antibiotic-free media were set as positive controls for growth. Detailed antibiotic information is provided in Table S8.

The detailed procedure was described as follows: briefly, approximately 2.5 g of sediments were mixed with 25 mL of a sterile water, the mixtures were stood for 20 min to enable the settling of large soil particles. Supernatants (approximately 100 μL) were transferred into 2216E medium supplemented with various antibiotics and incubated at 28 °C for 3-4 days. Resistant bacterial colonies that grew on the media were then harvested and washed with PBS (pH 7.4), and sent for metagenomic sequencing, following the “DNA Extraction and Metagenomic Sequencing” procedures.

Bacterial isolates obtained from PB, TE, CAZ, STR, and NA antibiotic-containing plates were subjected to antimicrobial susceptibility testing using the Kirby-Bauer disk diffusion method, the amount of each antibiotic disc (Liofilchem, Italy): PB (300 units), TE (30 µg), CAZ (30 µg), STR (10 µg), and NA (30 μg). The strains of *Escherichia coli* ATCC 25922 was used as the quality control.

**Alpha and Beta Diversity Analysis**

Shannon, pielou and richness indices for characterizing Alpha diversity of antibiotic-resistant genes were calculated using the vegan package (v2.6.8) in R software. Beta diversity of community composition was assessed through Non-metric Multidimensional Scaling (NMDS), based on Bray-Curtis distance matrices generated by the metaMDS function in vegan. Differences among all groups, as well as pairwise community comparisons, were evaluated using Permutational Multivariate Analysis of Variance (PERMANOVA) using the adonis2 function in vegan package (v2.6.8), based on 999 permutations of Bray–Curtis dissimilarities. All results were visualized using ggplot2 (v3.5.0).

**Risk ranking of Antibiotic Resistance Genes**

The risk-ranking framework for ARGs is adapted from established methodologies , employing a decision-tree-based classification system that stratifies ARGs into four risk ranks (Rank I–IV) based on three key criteria: (1) enrichment in human-associated environments, (2) mobility (mediated by MGEs), and (3) association with pathogenic hosts. Specifically, ARGs exhibiting <100-fold lower abundance in human-associated environments (e.g., human/animal feces, sewage, and contaminated sites) relative to non-impacted environments (e.g., marine water and natural soil) are categorized as ‘Rank IV’ (not human-associated). Among the human-associated ARGs, those confirmed as non-mobile through screening against the MGEs database are assigned to ‘Rank III’ (human associated and non-mobile). Human-associated ARGs that are mobile but lack documented associations with pathogenic hosts are classified as ‘Rank II’ (future threats). Finally, ARGs fulfilling all three criteria - (i) enriched in human-associated environments, (ii) mobile (MGE-associated), and (iii) carried by known pathogens - are designated as ‘Rank I’ (current threats), representing the highest risk tier.

**Metagenomic assembly for identification and abundance profiling of MGEs**

DIAMOND BLASTP (v2.16.0+) was used to search protein sequences derived from ORFs predicted in contigs against the MGE reference database mobileOG-db . An ORF was considered MGE-like if its alignment to a reference sequence exhibited an E-value ≤ 1e^-7^, similarity of ≥ 60% and query coverage ≥ 60%. We annotated the ARGs and MGEs on all contigs and then identified the contigs carrying both ARGs and MGEs, ARGs with potential mobility were defined as sharing a nearby area (< 10 kb) . The genetic structure plot was visualized using ggplot2 (v3.5.0).

The abundance of MGEs was quantified using Reads Per Kilobase per Million mapped reads (RPKM). RPKM values were calculated with CoverM (v0.7.0) based on BAM files generated from mapping sequencing reads to MGE regions. The following parameters were applied: --no-zeros, --min-read-percent-identity 95, and --min-read-aligned-percent 75.

**Source tracking of Antibiotic Resistance Genes**

To infer the probable sources of ARGs in hadal sediments, a Bayesian-based machine learning classifier, SourceTracker (v2.0.1) , was employed. This approach leverages a matrix of ARG subtypes and their relative abundances to predict source contributions, following established protocols . The model construction and ARG source prediction workflow commenced with the acquisition of 594 metagenomic datasets spanning four ecotypes: human feces (HF), animal feces (AF), wastewater treatment plants (WA), and natural environments (NT). These datasets were retrieved from public repositories, including the NCBI Sequence Read Archive (NCBI-SRA; https://www.ncbi.nlm.nih.gov/sra/), MG-RAST; http://metagenomics.anl.gov/), and the Human Microbiome Project Data Analysis and Coordination Center (HMPDACC; <http://hmpdacc.org/HMIWGS/all/>). Raw sequencing data for each ecotype were subjected to quality control and preprocessing using Trimmomatic (v0.38) with parameters “LEADING:2, TRAILING:2, SLIDINGWINDOW:4:15, HEADCROP:5, MINLEN:50, AVGQUAL:20” to ensure high-quality reads. Subsequently, ARGs within these datasets were identified and quantified using the ARGs-OAP pipeline . The resulting ARG subtype abundance matrices - comprising subtype classifications and their corresponding relative abundances - served as the training dataset for constructing the “source” models. SourceTracker was configured with optimized parameters (--sink_rarefaction_depth 0 --source_rarefaction_depth 0 --beta 1 --alpha1 0.001 --alpha2 0.001) to build the source models and predict the potential contributions of the four ecotypes to ARGs in hadal trench sediments. To mitigate false positive predictions , each SourceTracker analysis was independently replicated three times.

**Phylogenetic analysis of MAGs and genes**

Phylogenetic analysis of MAGs from hadal trench sediments was conducted based on 120 single-copy marker genes predicated and concatenated by GTDB-tk . The marker gene sequences were aligned using MAFFT (v7.475) and trimmed using trimAl (v1.4) with the automated1 setting. The maximum-likelihood (ML) phylogenomic tree was constructed using the concatenated aligned marker gene sequences with RAxML (v. 8.2.12) with 1000 bootstraps.

We performed phylogenetic analyses on four genes of interest, *fosX*, *sul2*, *arnA*, and *abeS* (Fig. S9) from metagenomic contigs assembled from sediments of six hadal trenches with candidates identified using DIAMOND (see “Metagenomic Assembly for Identification of ARG-like Open Reading Frames”). Published reference proteins encoded by these genes were downloaded from NCBI Identical Protein Groups database. NCBI reference proteins were clustered with CD-HIT (v4.8.1) at 99% identity for *fosX*, *sul2*, and *abeS* (parameters: -c 0.99 -G 0 -aS 0.9 -g 1 -d 0) and at 95% identity for *arnA* (parameters: -c 0.95 -G 0 -aS 0.9 -g 1 -d 0). The clustered references and hadal sequences were aligned with MAFFT (v7.475) , and trimmed using trimAl (v1.4) with the automated1 setting. The ML trees for the proteins were constructed by RAxML (v. 8.2.12) with settings of “-x 12345 -p 12345 -f a -m PROTGAMMALG -# 100”.

All phylogenetic trees were visualized using the interactive Tree Of Life (iTOL v6) tool . For clearer visualization of the phylogenetic relationships, the tree was midpoint-rooted using “root the tree at the midpoint” function in iTOL.

**Statistical analysis and presentation**

The statistical significance of ARG abundance among different years of Mariana Trench was compared using the One-way ANOVA test, followed by Tukey’s multiple comparison test with Tukey’s HSD for post-hoc analysis. The statistical significance of ARG abundance among different hadal trenches in Fig. 2a and the alpha diversity (Shannon, Richness and Pielou indexs) of ARG subtypes in Fig. S5, were compared using the nonparametric Kruskal-Wallis test, followed by Dunn’s multiple comparison test with Bonferroni correction for post-hoc analysis. All statistical analyses were performed in GraphPad Prism(v10.4.2). Column graphs were plotted in GraphPad Prism (v10.4.2).

**Supplementary Figure**

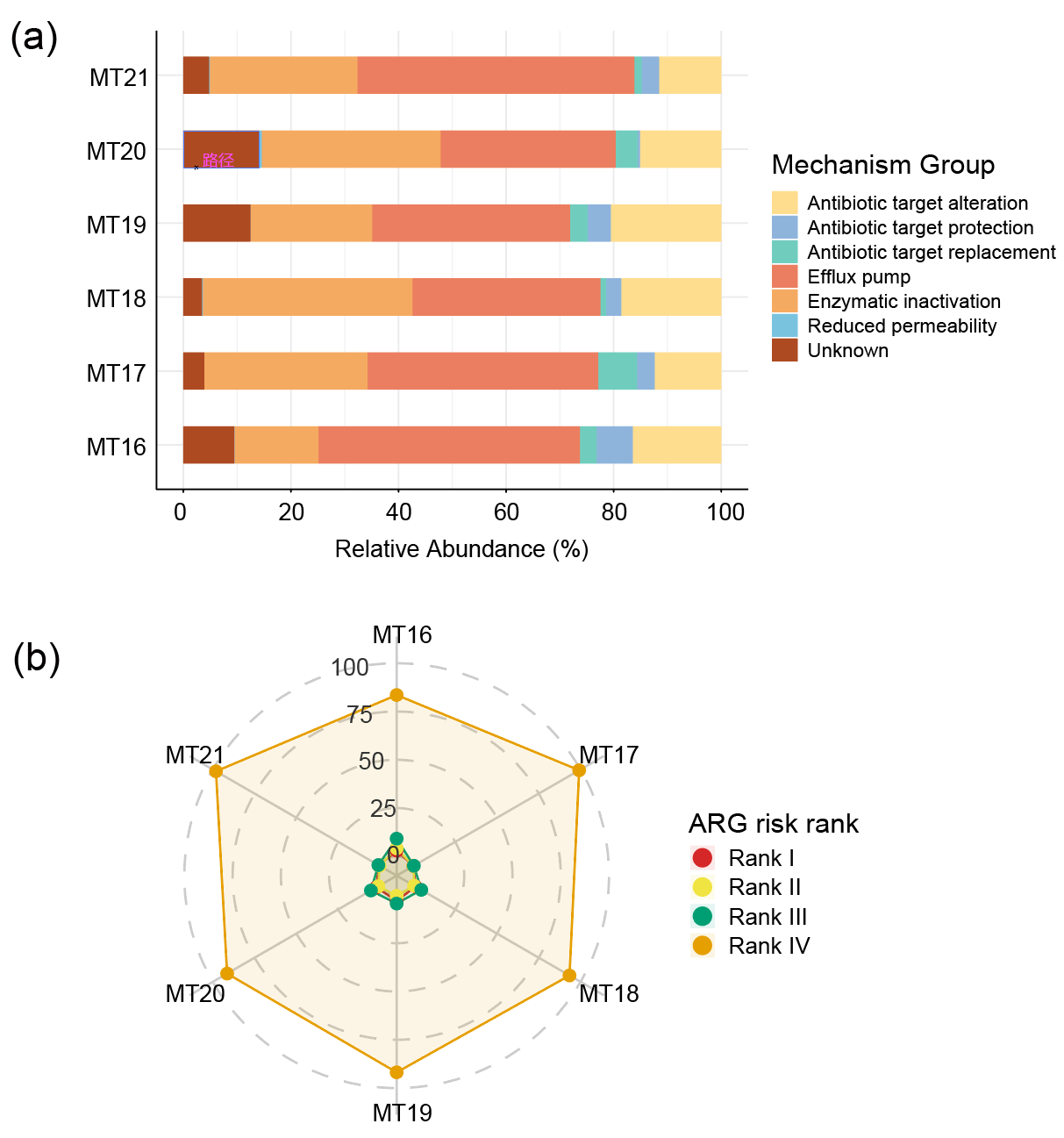

**Fig. S1 | Dynamics of antibiotic resistance mechanisms and risk rank profiles in Mariana Trench Challenger Deep sediments over the six-year period.** (a) Composition of ARG resistance mechanisms in Mariana Trench Challenger Deep sediments over the six-year study period. (b) Radar plot showing the relative proportion of different risk ranks of ARGs distributed in the Challenger Deep sediments of Mariana Trench over the six-year study period.

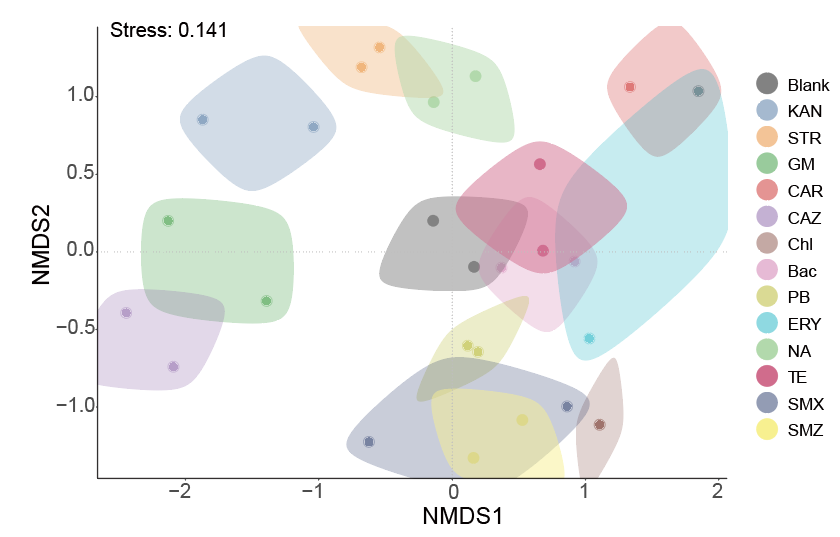

**Fig. S2 | NMDS analysis based on Bray-Curtis distance showing the overall microbial community structure at the genus level (stress = 0.141)**. The statistical significance of differences between groups was tested by PERMANOVA (p = 0.001, R² = 0.846).

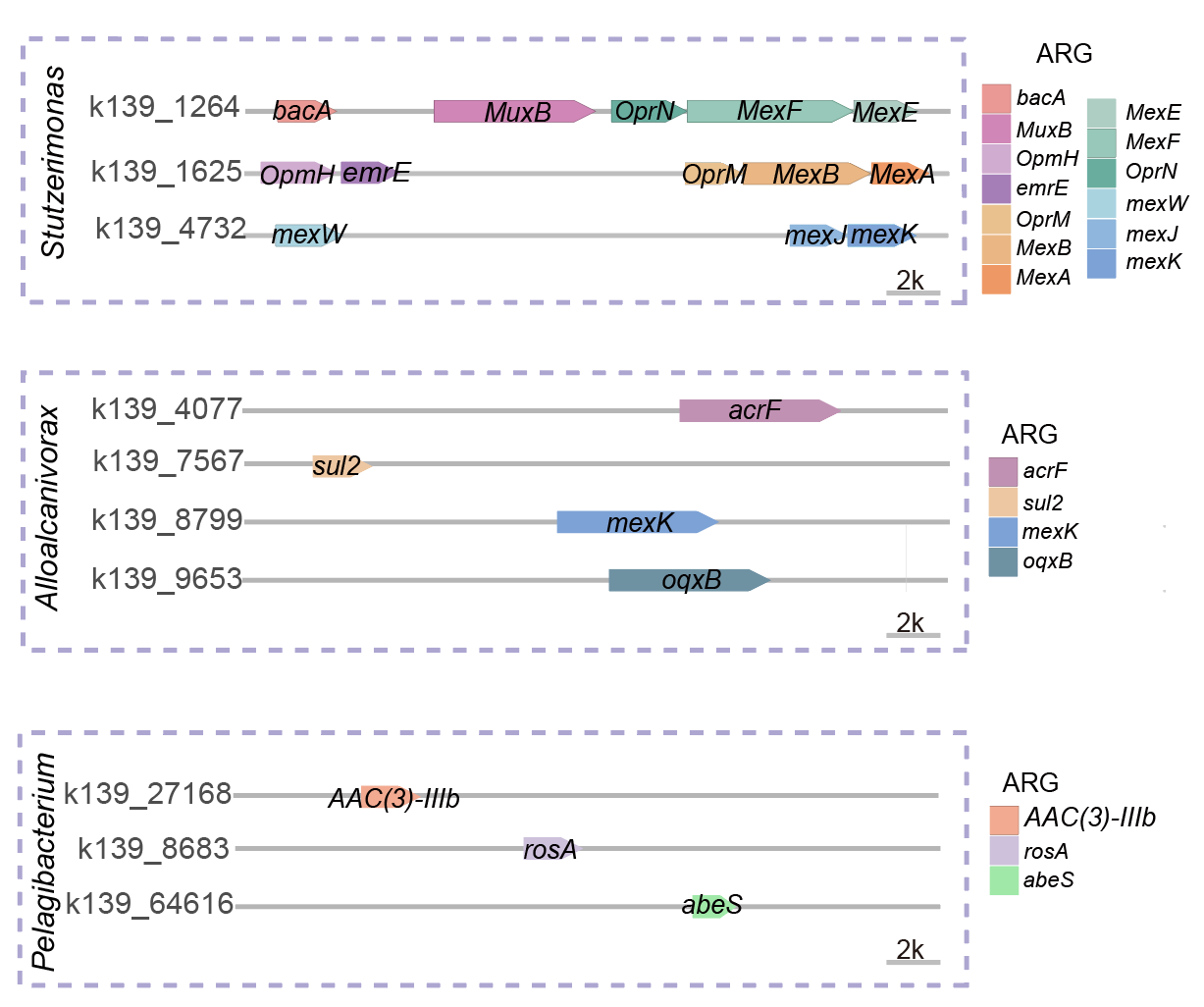

**Fig. S3 | Correspondence between phenotypic resistance and genotypes in bacteria selected on antibiotic-containing plates.**

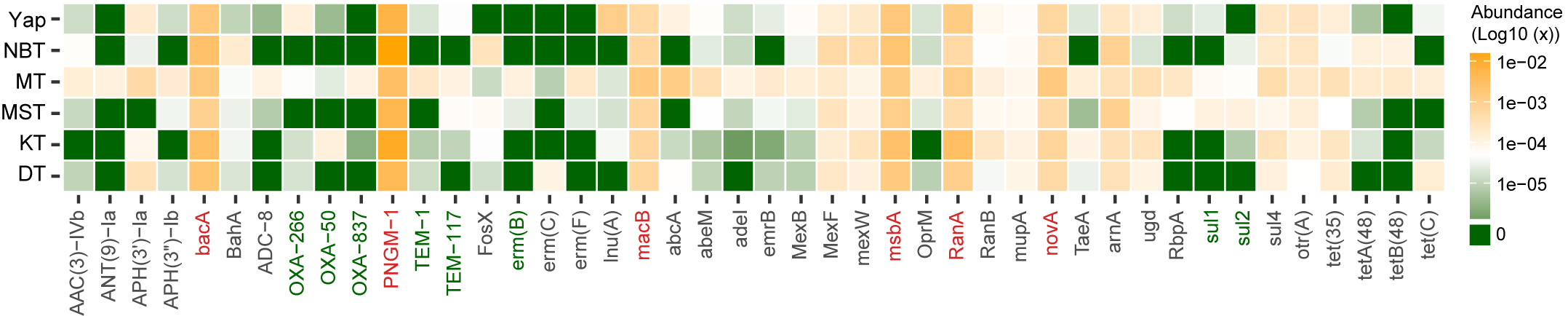

**Fig. S4 | Heatmap of the relative abundance of some ARG subtypes detected in the six hadal trenches.** Detailed abundance for all ARG subtypes were provided in Table S15. Abundance was log-transformed (i.e., Log10 (copy/cell)) for the plot. MLS: macrolide, lincosamide, and streptogramin.

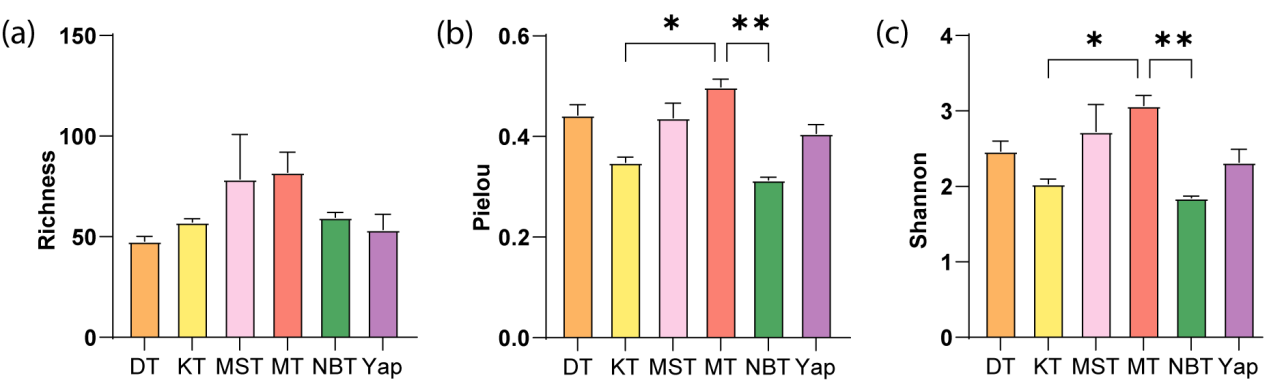

**Fig. S5 | Alpha diversity indexs of ARGs in the six hadal trenches.** (a) richness index; (b) pielou’s evenness index; (c) shannon index; Each error bar corresponds to the SEM and data were showed as mean + SEM. Dunn’s multiple comparison test with Bonferroni correction was performed to obtain the *p* value. *p<0.05; **p<0.01. Non-significant differences between trenches in some alpha diversity indexes were not labeled. MT: Mariana Trench; NBT: New Britain Trench; Yap: Yap Trench; MST: Massau Trench; KT: Kermadec Trench.

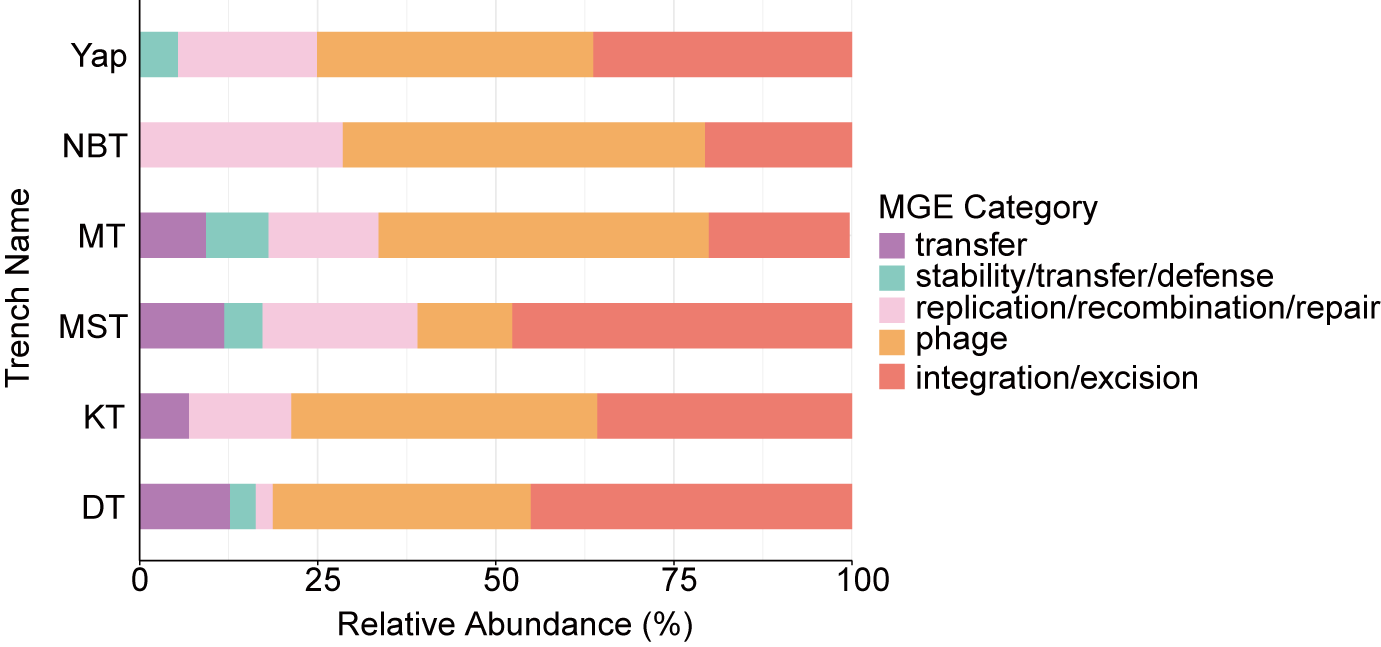

**Fig. S6 | Bar plot of the distribution of MGE categories in the six hadal trench sediments.** MT: Mariana Trench; NBT: New Britain Trench; Yap: Yap Trench; MST: Massau Trench; KT: Kermadec Trench.

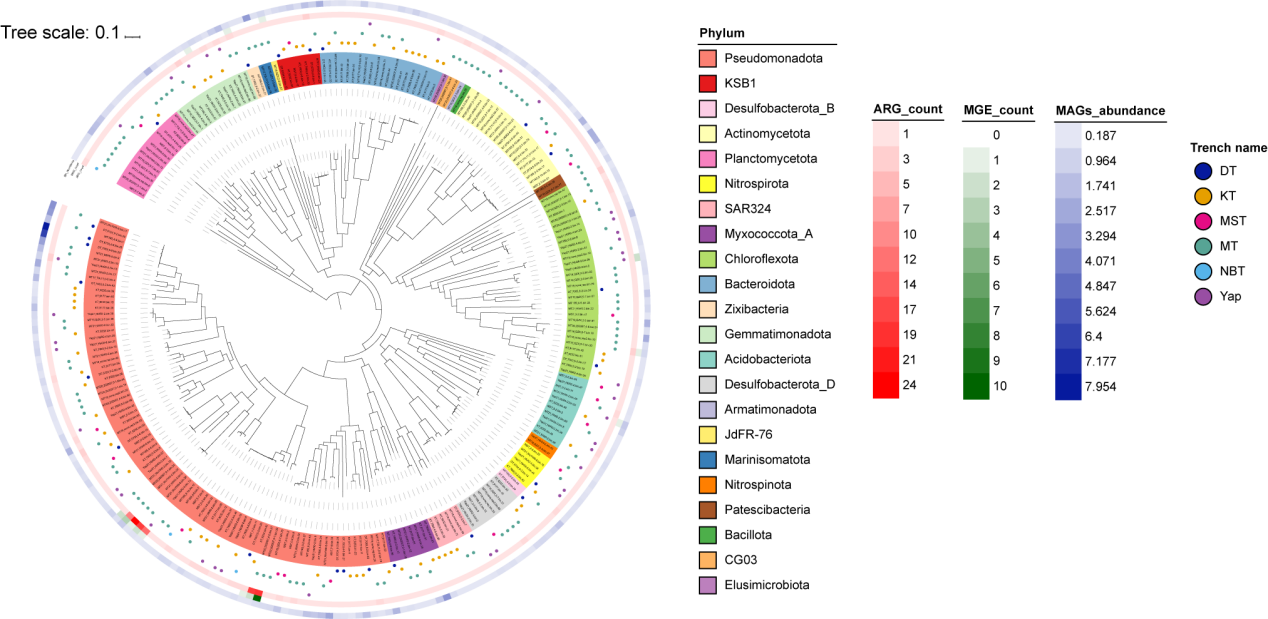
**Fig. S7 | Maximum-likelihood phylogenetic tree of the ARG-carrying MAGs in the six hadal trench sediments.** The colors of the strip of outer layer show the taxonomy at the phylum level. The blue, yellow, red, green, cyan and purple dots present MAGs belong to Diamantina Trench (DT), Massau Trench (MST), Mariana Trench (MT), New Britain Trench (NBT), Yap Trench (Yap), and Kermadec Trench (KT). Purple bars show the prevalence of the MAGs in the six hadal trench sediments. Red bars indicates the number of ARGs in the MAGs. Green bar indicates the number of MGEs in the MAGs.

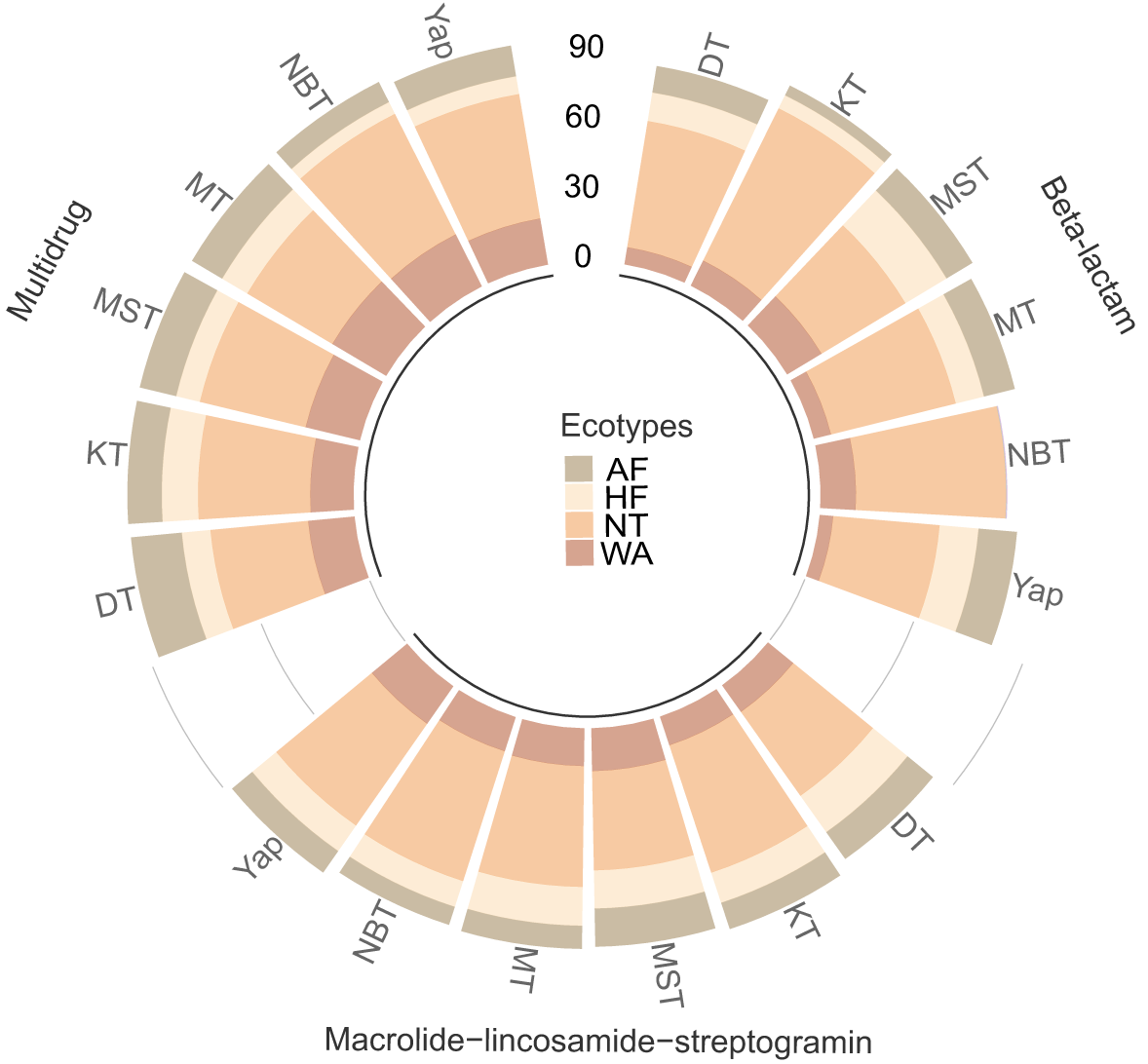

**Fig. S8 | Relative proportion of predicted sources for the three dominant ARG types, i.e., multidrug, macrolide-lincosamide-streptogramin and beta-lactam, detected in the hadal trench sediments.** NT: natural environment; AF: animal feces; HF: human feces; WA: wastewater treatment plant; MT: Mariana Trench; NBT: New Britain Trench; Yap: Yap Trench; MST: Massau Trench; KT: Kermadec Trench.

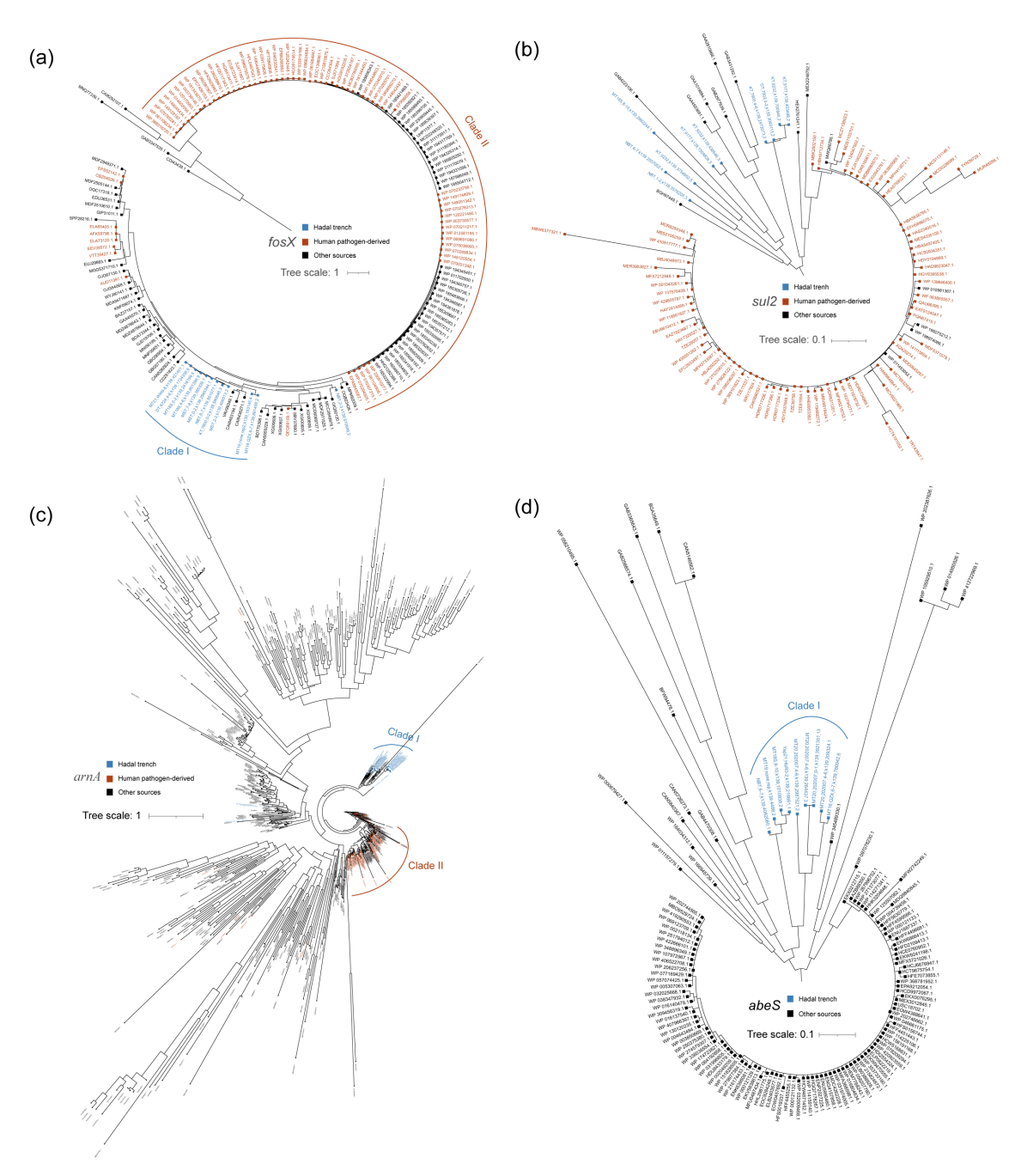

**Fig. S9 | Phylogenetic analysis of four ARGs identified in hadal environments and their homologous sequences: *fosX* (a), *sul2* (b), *arnA* (c), and *abeS* (d).** Sequences derived from hadal trench samples are highlighted in blue. Homologous sequences retrieved from the NCBI database are annotated with their accession numbers on corresponding tree leaves; those originating from human pathogens are colored red, while sequences from other sources are shown in black. Environmental sources for selected reference sequences, listed by gene and panel: a, *fosX*: CAI8463194.1 & CAI8436271.1 (North Pacific Ocean), VAV99349.1 (hydrothermal vent metagenomes); b, *sul2*: GAB4223106.1 (hot spring), BGH97449.1 (marine), HEX2761241.1 (USA wetland soil), MEX2248752.1 (Ancient Andean lake sediments); c, *arnA*: WP_136062485.1 (Black Sea sediment), WP_228854127.1 (deep-sea hydrothermal vent chimney), GBE13456.1 & GBE15225.1 (sub-seafloor sulfide deposits). Note: for *abeS*, all homologous sequences including those from human pathogens are depicted in black.

**Supplemental Tables**

**Tables S1 to S25:**

**Table S1.** Sample information of hadal trench sediments.

**Table S2.** The normalized abundance of the total ARGs in Mariana Trench Challenger Deep sediments over the six-year study period.

**Table S3.** Abundance of ARG types in Mariana Trench Challenger Deep sediments over the six-year study period.

**Table S4.** Abundance of ARG subtypes in Mariana Trench Challenger Deep sediments over the six-year study period.

**Table S5.** Relative abundance of ARGs the different type of resistance mechanisms in Mariana Trench Challenger Deep sediment samples over the six-year study period.

**Table S6.** Relative Abundance of ARGs with efflux pump mechanism detected in the Challenger Deep sediments of Mariana Trench over the six-year study period.

**Table S7.** Relative abundance of different ranks of ARGs detected in the Challenger Deep sediments of Mariana Trench over the six-year study period.

**Table S8.** Antibiotic categories and concentrations.

**Table S9.** The information of the metagenomic datasets and quality statistics of metagenomic assembly.

**Table S10.** MAGs recovered from different treatments after quality assessment (completeness>50% and contamination<10%).

**Table S11.** The information of contigs carrying one ARG as well as the diversity and distribution of ARGs in each sample.

**Table S12.** Summary of the ARG-carrying MAGs recovered from different treatments.

**Table S13.** The normalized abundance of the total ARGs in the six hadal trench sediments.

**Table S14.** Abundance of ARG types detected in the six hadal trench sediment samples.

**Table S15.** Abundance of ARG subtypes in the six hadal trench samples (copy/cell).

**Table S16.** Relative abundance of ARGs with different resistance mechanisms in the six hadal trench samples.

**Table S17.** Relative abundance of different ranks of ARGs detected in the six hadal trench samples.

**Table S18.** Summary of the metagenomic datasets and quality statistics of metagenomic assembly for hadal sediment samples.

**Table S19.** Summary of the ARG-carrying contigs recovered from six hadal trench samples.

**Table S20.** Co-occurrence of ARGs and MGEs in contigs.

**Table S21.** MAGs recovered from different trench samples after quality assessment (completeness>50% and contamination<10%) and dereplication.

**Table S22.** Summary of the ARG-carrying MAGs recovered from six hadal trench samples.

**Table S23.** Information of 594 collected environmental datasets for source prediction.

**Table S24.** Relative contribution of various sources (i.e. AF, HF, NT and WA) to the ARGs in the six hadal trench predected by SourceTracker.

**Table S25.** Relative contribution of various sources (i.e. AF, HF, NT and WA) to three dominant ARG types in the six hadal trench predected by SourceTracker.

**References**

[1]	Peoples LM, Grammatopoulou E, Pombrol M, et al. Microbial community diversity within sediments from two geographically separated hadal trenches. Front Microbiol, 2019, 10: 347

[2]	Li Y, Liu H, Xiao Y, et al. Metagenome sequencing and 982 microbial genomes from kermadec and diamantina trenches sediments. Sci Data, 2024, 11: 1067

[3]	Zhang W, Li Y, Chu Y, et al. Deep-sea ecosystems as an unexpected source of antibiotic resistance genes. Mar Drugs, 2024, 23: 17

[4]	Zhang X, Xu W, Liu Y, et al. Metagenomics reveals microbial diversity and metabolic potentials of seawater and surface sediment from a hadal biosphere at the yap trench. Front Microbiol, 2018, 9: 2402

[5]	Xiao X, Zhao W, Song Z, et al. Microbial ecosystems and ecological driving forces in the deepest ocean sediments. Cell, 2025, 188: 1363-1377

[6]	Zhou YL, Mara P, Cui GJ, et al. Microbiomes in the challenger deep slope and bottom-axis sediments. Nat Commun, 2022, 13: 1515

[7]	Zhang RY, Wang YR, Liu RL, et al. Metagenomic characterization of a novel non-ammonia-oxidizing thaumarchaeota from hadal sediment. Microbiome, 2024, 12: 7

[8]	Cheng H, Zhang Y, Guo Z, et al. Microbial dimethylsulfoniopropionate cycling in deep sediment of the mariana trench. Appl Environ Microbiol, 2023, 89: e0025123

[9]	Minas K, McEwan NR, Newbold CJ, et al. Optimization of a high-throughput ctab-based protocol for the extraction of qpcr-grade DNA from rumen fluid, plant and bacterial pure cultures. FEMS Microbiol Lett, 2011, 325: 162-169

[10]	Bolger AM, Lohse M, Usadel B. Trimmomatic: A flexible trimmer for illumina sequence data. Bioinformatics, 2014, 30: 2114-2120

[11]	Shen W, Le S, Li Y, et al. Seqkit: A cross-platform and ultrafast toolkit for fasta/q file manipulation. PLoS One, 2016, 11: e0163962

[12]	Bushnell B. Bbmap: A fast, accurate, splice-aware aligner. 2014,

[13]	Li D, Luo R, Liu CM, et al. Megahit v1.0: A fast and scalable metagenome assembler driven by advanced methodologies and community practices. Methods, 2016, 102: 3-11

[14]	Uritskiy GV, DiRuggiero J, Taylor J. Metawrap-a flexible pipeline for genome-resolved metagenomic data analysis. Microbiome, 2018, 6: 158

[15]	Yin X, Zheng X, Li L, et al. Args-oap v3.0: Antibiotic-resistance gene database curation and analysis pipeline optimization. Engineering, 2023, 27: 234-241

[16]	Buchfink B, Reuter K, Drost HG. Sensitive protein alignments at tree-of-life scale using diamond. Nat Methods, 2021, 18: 366-368

[17]	von Meijenfeldt FAB, Arkhipova K, Cambuy DD, et al. Robust taxonomic classification of uncharted microbial sequences and bins with cat and bat. Genome Biol, 2019, 20: 217

[18]	Chaumeil PA, Mussig AJ, Hugenholtz P, et al. Gtdb-tk: A toolkit to classify genomes with the genome taxonomy database. Bioinformatics, 2019, 36: 1925-1927

[19]	Hyatt D, Chen GL, Locascio PF, et al. Prodigal: Prokaryotic gene recognition and translation initiation site identification. BMC Bioinformatics, 2010, 11: 119

[20]	Zhang AN, Gaston JM, Dai CL, et al. An omics-based framework for assessing the health risk of antimicrobial resistance genes. Nat Commun, 2021, 12: 4765

[21]	Brown CL, Mullet J, Hindi F, et al. Mobileog-db: A manually curated database of protein families mediating the life cycle of bacterial mobile genetic elements. Appl Environ Microbiol, 2022, 88: e0099122

[22]	Sun J, Liao XP, D'Souza AW, et al. Environmental remodeling of human gut microbiota and antibiotic resistome in livestock farms. Nat Commun, 2020, 11: 1427

[23]	Aroney STN, Newell RJP, Nissen JN, et al. Coverm: Read alignment statistics for metagenomics. Bioinformatics, 2025, 41: btaf147

[24]	Knights D, Kuczynski J, Charlson ES, et al. Bayesian community-wide culture-independent microbial source tracking. Nat Methods, 2011, 8: 761-763

[25]	Li LG, Yin X, Zhang T. Tracking antibiotic resistance gene pollution from different sources using machine-learning classification. Microbiome, 2018, 6: 93

[26]	Zhang Y, Zhang B, Ahmed I, et al. Profiles and natural drivers of antibiotic resistome in multiple environmental media in penguin-colonized area in antarctica. Fundam Res, 2025, 5: 269-281

[27]	Henry R, Schang C, Coutts S, et al. Into the deep: Evaluation of sourcetracker for assessment of faecal contamination of coastal waters. Water Res, 2016, 93: 242-253

[28]	Yamada KD, Tomii K, Katoh K. Application of the mafft sequence alignment program to large data-reexamination of the usefulness of chained guide trees. Bioinformatics, 2016, 32: 3246-3251

[29]	Capella-Gutiérrez S, Silla-Martínez JM, Gabaldón T. Trimal: A tool for automated alignment trimming in large-scale phylogenetic analyses. Bioinformatics, 2009, 25: 1972-1973

[30]	Stamatakis A. Raxml version 8: A tool for phylogenetic analysis and post-analysis of large phylogenies. Bioinformatics, 2014, 30: 1312-1313

[31]	Fu L, Niu B, Zhu Z, et al. Cd-hit: Accelerated for clustering the next-generation sequencing data. Bioinformatics, 2012, 28: 3150-3152

[32]	Letunic I, Bork P. Interactive tree of life (itol) v6: Recent updates to the phylogenetic tree display and annotation tool. Nucleic Acids Res, 2024, 52: W78-W82
